# Hypoxia alters the effects of hypomethylating agents in acute myeloid leukaemia cells

**DOI:** 10.1101/2023.12.07.570313

**Authors:** Sam Humphries, Sean M. Burnard, Simon Keely, Danielle R. Bond, Heather J. Lee

## Abstract

**Background:** Acute myeloid leukaemia (AML) is a deadly haematological malignancy that originates from mutated myeloid progenitor cells that lie quiescent in the hypoxic bone marrow. Elderly patients who cannot tolerate standard chemotherapies are administered low-dose hypomethylating agents (HMA) which act in a replication-dependent manner to reprogram the epigenome. Relapse is common following HMA treatment and may arise from quiescent leukaemia cells in the hypoxic bone marrow. Therefore, the effects of hypoxia on HMA efficacy may influence AML progression.

**Results:** AML cell lines (MOLM-13, MV-4-11, HL-60) were treated with decitabine (100nM) or azacitidine (500-2000nM) in normoxic (21% O_2_) and hypoxic (1% O_2_) conditions. Exposure to hypoxia significantly reduced AML cell growth across all cell lines, with no additional effects observed upon HMA treatment. This was associated with distinct effects on DNA methylation. The extent of hypomethylation induced by AZA treatment was reduced in hypoxia, whereas DAC-induced hypomethylation was maintained in low oxygen conditions. Transcriptional response to HMA treatment were also altered in hypoxia, with HMAs failing to up-regulate antigen presentation pathways in hypoxia. In particular, human leukocyte antigens (HLAs) such as HLA-DR were increased upon HMA treatment in normoxia, but not hypoxia.

**Conclusion:** Our results suggest that HMA-induced antigen presentation may be impaired in hypoxic tissues such as the bone marrow. This study highlights the need to consider microenvironmental factors when designing co-treatment strategies to improve HMA therapeutic efficacy.

## 1. Introduction

Acute myeloid leukaemia is a deadly haematological malignancy, with a 5-year survival rate of ∼26% [1-4]. Characterised by the enhanced proliferation and diminished differentiation of myeloid progenitor cells, AML predominantly develops in those over the age of 60 [4, 5]. While standard chemotherapies, like daunorubicin and cytarabine, are effective in most patients, relapse remains prevalent, with recurrence typically occurring within 3 years of diagnosis [6, 7]. Due to the cytotoxic nature of these chemotherapies and the harsher effects observed in elderly or immunocompromised patients, alternative low dose therapeutic regimens have been developed that can target epigenetic modifications, like DNA methylation.

DNA methylation is a crucial component of the epigenome, with important roles in regulating gene expression [8]. In AML, mutations in epigenetic modifiers have been identified in ∼20% of AML patients, often promoting a dysregulated transcriptional profile that encourages the AML phenotype [4, 9-15]. Given the vital role that DNA methylation plays in AML progression, hypomethylating agents (HMAs), like decitabine (DAC) and azacitidine (AZA), have been approved for clinical use against both pre-leukaemic myelodysplastic syndrome (MDS) and AML [16]. Mechanistically, HMAs are cytidine analogues that are incorporated into DNA during replication [17-19]. At low doses, HMAs have been shown to restructure the epigenome by trapping and degrading the activity of DNMT enzymes, promoting global DNA de-methylation. The resulting HMA-induced promoter hypomethylation has, in turn, been found to prompt the re-expression of tumour suppressor genes that can consequently decrease AML growth and increase immunogenicity [19-21]. While these results seem promising, only 30-50% of leukaemia patients respond to treatment [22-24], and the epigenetic and transcriptional changes induced by treatment do not provide a strong prediction of patient response. This suggests that other factors, such as components of the tumour microenvironment, may limit treatment efficacy [25].

Leukaemic stem cells (LSCs) reside within the hypoxic environment of the bone marrow, where oxygen tensions range from 1-5% O_2_ [26, 27]. Here, it is believed that LSCs adopt similar quiescent properties to healthy haematopoietic stem cells, allowing them to lie quiescent and protected away from extracellular stressors, like reactive oxygen species, that can induce DNA damage [4]. It is also believed that these quiescent properties allow LSCs to avoid chemotherapies, promoting chemoresistance and, in turn, relapse [4, 28-34]. This has been demonstrated in MDS patients, in which non-responding patients treated with AZA showed a higher proportion of quiescent (G0) progenitor cells within the BM [35]. Unfortunately, insight into the effect of this hypoxic microenvironment on HMA efficacy in AML is lacking. Our study therefore aims to investigate the direct effects of hypoxia on AML cell activity, and whether this, in turn, imposes implications for HMA efficacy and subsequent downstream transcriptional activity.

## 2. Methods

### Routine cell culture

AML cell lines, devoid of mutations in key epigenetic regulators, were routinely cultured at 5×10^5^ cells/mL in normoxic conditions (21% O_2_, 5% CO_2_, 37°C), until cell viability was stable >90%. MOLM-13 and MV-4-11 cell lines were cultured using RPMI-1640 medium (Sigma-Aldrich, U.S.) supplemented with 2mM GlutaMAX (Thermofisher, U.S.) and 10% Foetal Bovine Serum (Sigma-Aldrich, U.S.), while HL-60s were cultured in Iscove’s Modified Dulbecco’s Medium (IMDM) (Sigma-Aldrich, U.S.) with 4mM GlutaMAX and 10% FBS. Cell lines were also tested for mycoplasma using the MycoAlert Mycoplasma detection kit (Lonza, Switzerland).

### HMA Treatment and Cell Growth analysis

After ensuring stable viability, AML cell lines were pre-exposed to either normoxic (21% O_2_, 5% CO_2_, 37°C) or hypoxic (1% O_2_, 5% CO_2_, 37°C) conditions for 72 hours, with appropriate medium pre-warmed and oxygen-equilibrated before use. A standard CO_2_ incubator was used for culture at 21% O_2_, while 1% O_2_ was maintained in a Hypoxic Glove-box Chamber (PLAS labs, U.S.). Following pre-exposure, cells were re-seeded into respective oxygen tensions at 4×10^5^ cells/mL and treated for a further 72 hours with Decitabine (DAC; 100nM for all cell lines) (Sellekchem, U.S.) or 5-Azacitidine (AZA; 500nM for MOLM-13 and MV-4-11 cells, and 2000nM for HL-60 cells) (Sellekchem, U.S.). Doses were selected based on previous dose response experiments that showed maximal loss of DNA methylation with minimal effects on cell viability (data not shown). Cell growth and viability were analysed using a trypan blue exclusion assay (Biotools, AUS), and 2×10^6^ cells were stored in 1X DNA/RNA Shield (Zymo Research, U.S.) at -80°C for downstream molecular analysis.

### Flow Cytometry

To track the cell division rate during HMA treatment in each oxygen tension, CellTrace Far Red Cell Proliferation Stain (Thermofisher, USA) was used. Prior to HMA exposure, cell lines were stained according to the manufacturer’s instructions, and a day 0 positive and unstained negative control were recorded using flow cytometry. CellTrace stained cells were then treated with HMAs for 72 hours under each oxygen tension, and cell division for each treatment was assessed based on decreases found in CellTrace stain using the following equation:

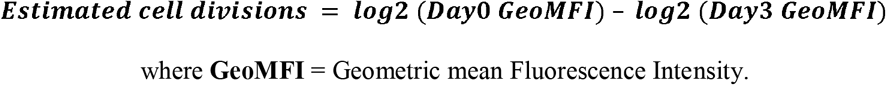

Human leukocyte antigen (HLA) surface marker expression was also investigated. At treatment endpoint, cells were stained with monoclonal antibodies targeting HLA-DR or -G, or a polyclonal antibody targeting HLA-DR, -DP and -DQ (***Supplementary Table 1***). Isotype control antibodies were also used to control for non-specific binding background signal of the HLA antibodies.

All cell staining was analysed using the FACSCanto II flow cytometer. Gating strategies were applied accordingly to remove any cell doublets and fluorescent signal was analysed using Kaluza software (Beckman Coulter Life Sciences, U.S.) (***Supplementary Figure 1***).

### Nucleic Acid Isolation & Purification

Nucleic acid extractions were performed on HMA-treated cells to analyse downstream DNA methylation and transcriptional changes. Using the Quick DNA/RNA Magbead Kit (Zymo, USA), both DNA and RNA were purified following the manufacturer’s instructions and eluted in UltraPure distilled water (Thermofisher, U.S) for quantification. DNA was quantified using the Qubit Broad Range dsDNA Assay kit (ThermoFisher, U.S.) or 1X High Sensitivity dsDNA assay kit (ThermoFisher, USA), and assessed by the Qubit® 2.0 Fluorometer. Alternatively, RNA was quantified using an Epoch plate reader at 260 & 280nm absorbances.

### Library Preparation for Bisulphite Sequencing

To analyse DNA methylation, sequencing libraries were prepared using a post-bisulphite adaptor tagging (PBAT) strategy as described [36]. Briefly, DNA purified from treated cells was first subjected to bisulphite conversion using the EZ-96 DNA Methylation-Direct MagPrep Kit (Zymo Research, U.S.; #D5044) to convert any unmodified cytosine residues to uracil. Samples were then resuspended in a pre-amplification mix: 1X blue buffer, 0.4mM dNTP mix, 2uM 6NF preamplification oligo (5’-CTACACGACGCTCTTCCGATCTNNNNNN-3’), and UltraPure distilled water, and incubated at 65°C to denature the DNA. Klenow (3’ → 5’ exo-, 50U) (Custom Science, AUS) was added to each sample and first strand synthesis was performed on a thermocycler by incubating as follows: 5min at 4°C, 37 repeats of +1°C every 15s, 30min at 37°C. Samples were then incubated with Exonuclease I (40U) (New England Biolabs, U.S) at 37°C for 1 hour and purified using Agencourt AMPure XP beads (0.8X) (Beckman Coulter, U.S.). For second strand synthesis, samples and beads were resuspended in a second-strand mixture, as indicated: 1X blue buffer, 0.4mM dNTP mix, 2uM 6NR_NEB adaptor 2 oligo (5’-CAGACGTGTGCTCTTCCGATCTNNNNNN-3’) and UltraPure distilled water and incubated at 95°C for 45s. Like first strand synthesis, Klenow was added to each sample and incubated on the thermocycler using the same conditions, before a second round of purification with AMPure XP beads (0.8X).

To amplify the libraries for sequencing, samples and beads were resuspended in KAPA HiFi HotStart ReadyMix (1X) (Millenium Science, AUS), which contains a DNA polymerase to amplify the DNA. Following the manufacturer’s instructions, specific i7 and i5 NEBNext index pairs (5uM) (New England Biolabs, U.S) were added to each sample and incubated as follows: 1 cycle at 95°C for 2min; 6 cycles of 94°C for 80s, 65°C for 30s and 72°C for 30s; 72°C for 3min; hold at 4°C. Amplified libraries were purified using AMPure XP beads (0.8X) and eluted in UltraPure distilled water. The quality of each sample library was assessed by analysing the fragment size distribution on an Agilent 4200 Tapestation system (Agilent, USA). A Qubit 2.0™ fluorometer was then used to quantify the concentration of amplified libraries, ready for sample normalisation, pooling and DNA sequencing.

### DNA Sequencing and DNA methylation Analysis

Sequencing was performed using a MiSeq Reagent Kit v2 (Illumina, U.S.) in 150bp paired-end mode, with a loading sample concentration of 10pM and a 5% PhiX (Illumina, U.S.) spike-in. Raw reads were trimmed in pair-end mode using Trim Galore (v0.6.5, Cutadapt v2.10, options ‘--clip_R1 9 -- clip_R2 9) to exclude poor-quality calls and adaptors [37, 38]. Alignment and methylation calling of trimmed reads was performed using Bismark (v0.22.3) in PBAT mode against the human GRCh38 reference assembly using a custom approach [39, 40] in which mapping is first performed in pair-end mode with unmapped singleton reads written out, followed by alignment of the singleton reads in single-end mode. The following parameters were used: i) -- unmapped –pbat -1 <read1> -2 <read2>, ii) --pbat --single_end <unmapped_read1> iii) --pbat --single_end <unmapped_read2>. Mapped reads were then deduplicated (using deduplcation_bismark) and methylation extraction performed (using bismark_methylation_extractor) in paired end and single end mode separately, and then merged into a single coverage file per sample. The methylation extraction reports obtained from Bismark were then used to determine the average global DNA methylation level of each sample.

### Transcriptome Analysis using RNA-Sequencing

RNA sequencing was performed by the Ramaciotti Centre for Genomics (UNSW, Australia) using the RNA extracted following HMA treatments in MOLM-13 cells. For each treatment condition, triplicate libraries were prepared using the Illumina stranded mRNA prep kit (Illumina, US) and sequencing was carried out with a NovaSeq 6000 SP 2×100bp FlowCell plus PhiX spike-in (Illumina, US), outputting ∼25M read pairs/sample. Raw reads were trimmed in pair-end mode using Trim Galore (v0.6.5, Cutadapt v2.10) in paired-end mode with default parameters and retaining unpaired reads. Hisat2 (v2.1.0) [41] was used to map and align trimmed and unpaired reads using default parameters (--phred33) to the human reference genome build (GRCh38), and samtools (v1.10) [42] was used to generate bam files. SeqMonk (v1.47.1), was used to perform quality control assessment using the ‘QC RNA plot’, ‘Distribution Plot’ and ‘DataStore Tree plot’ functions.

To determine genes that were differentially expressed between treatment groups, DESeq2 analysis [43] was performed in SeqMonk. Specifically, differential expressed genes (probes) were generated by comparing untreated samples with HMA treatment groups within respective oxygen tensions. Genes that had a significant change (p<0.01, |log_2_(fold change)| > 1) in expression for each analysis were then assembled into appropriate probe ‘lists’. Probe lists for each comparison were then compiled to form a heatmap using the per-probe normalized ‘Hierarchical clustering’ function, allowing for visualization of common or differential gene expression trends between untreated samples and HMA treatments in both normoxia and hypoxia conditions.

Overlap of genes up- (log_2_(fold change > 1) or down-regulated (log_2_(fold change < -1) by HMA treatment in 21% or 1% O_2_ were then displayed as a venn diagram using R (ggvenndiagram [44]), isolating genes uniquely up- or down-regulated in these conditions. Gene ontology over representation analysis (biological process) was performed on the unique gene sets by ClusterProfiler [45] using compareCluster(fun=“enrichGo”, ont=“BP”, padj=“fdr”) and displayed using enrichplot::treeplot() [46] for the top 15 terms with default parameters.

## 3. Results

### Hypoxia decreases AML cell growth irrespective of HMA treatment

To explore the impact of hypoxia on HMA efficacy in AML, we first assessed the effects on cell growth, viability, and cell division. To do this, AML cell lines (MOLM-13, MV-4-11, HL-60) were cultured in 21% or 1% O_2_ and treated with decitabine (2-deoxy-5-azacytidine, DAC) or azacytidine (AZA) at respective doses for 72 hours. In normoxic conditions (21% O_2_), DAC significantly reduced cell counts relative to untreated controls in all cell lines, while AZA was most notably effective in HL-60 cells (***Fig. 1A***). It is worth noting here however, that the observed decrease in HL-60 growth is most likely attributed to the higher dose (2000nM) used compared to MOLM-13 and MV-4-11 (500nM) for the maximal reduction of DNA methylation. When introducing hypoxia (1% O_2_), cell growth was dramatically reduced in all treatment conditions, with comparable counts observed between HMA-treated and untreated samples. While significant decreases in cell viability were observed in some conditions (e.g., in untreated MOLM-13 cells; p<0.005) (***Fig. 1B***), they were insufficient to explain the overall reductions in viable cells observed after 72 hours of treatment in hypoxia.

**Figure 1:**
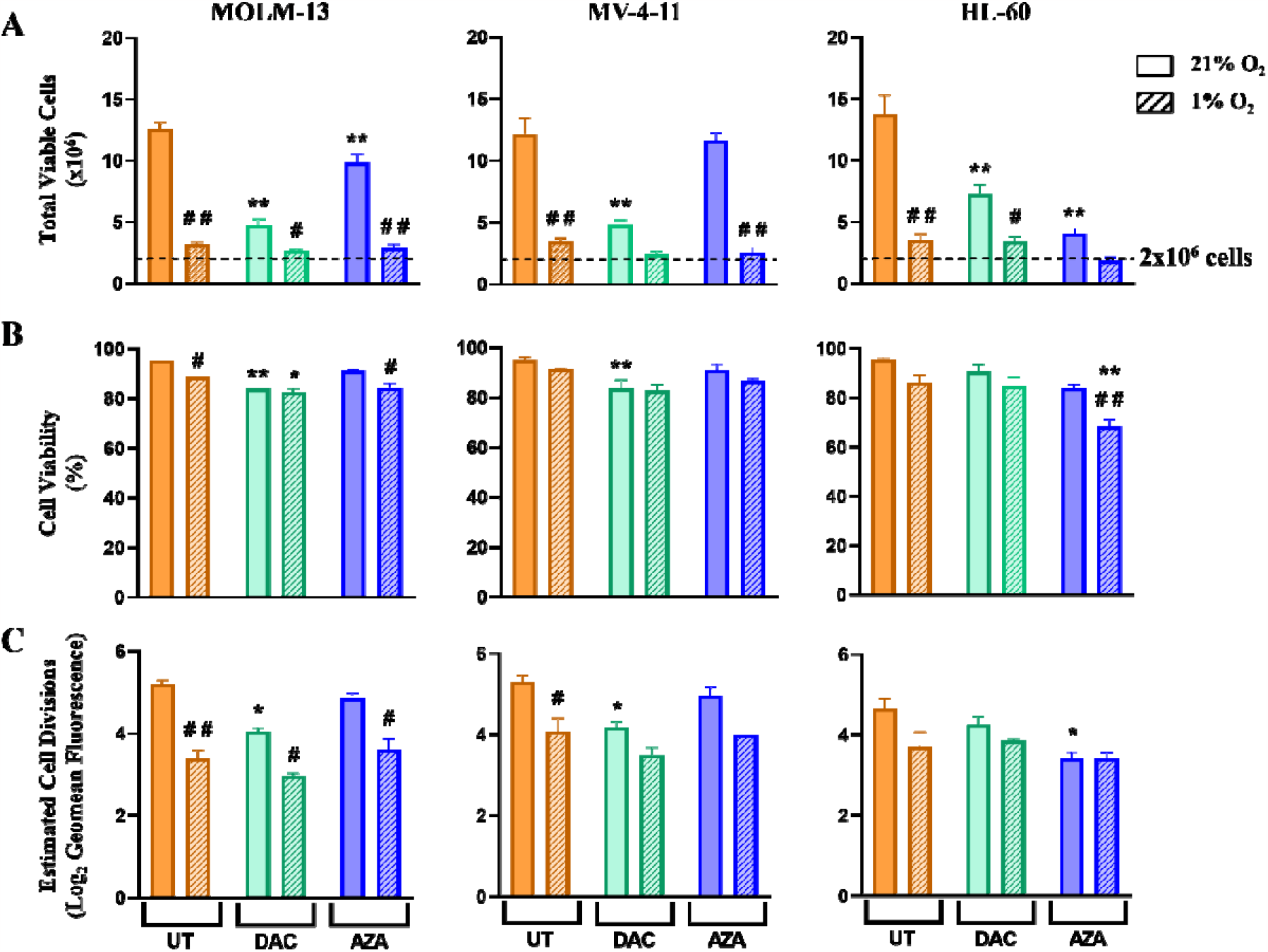
*The effect of hypoxia and HMA treatment on AML cell growth, viability, and cell division*. AML cell lines (MOLM-13, MV-4-11, HL-60) were first pre-conditioned to either normoxic (21% O_2_; solid) and hypoxic (1% O_2_; striped) conditions for 72 hours. Following pre-exposure, cells were re-seeded at 4×10^5^ cells/mL (A; dashed line) in respective oxygen tensions and treated with either DAC (100nM) or AZA (500-2000nM) for a subsequent 72 hours. Trypan blue exclusion assays were used to detect: (**A**) the total number of live cells (trypan blue negative cells), and (**B**) the percentage of viable cells. The number of cell divisions occurring during treatment was also estimated using CellTrace fluorescence. (**C**) shows the CellTrace fluorescence geomean transformed to a delta log_2_ expression to represent a theoretical number of cell divisions over the 72-hour time course with and without treatment. Data is presented as mean ± standard error of the mean for replicate experiments (MOLM-13 n=10; MV-4-11 n= 6; HL-60 n=6). # indicates significant difference between oxygen tensions within each treatment group. * indicates significant difference of treatment from UT control within each oxygen tension. Two-way ANOVA (Tukey’s Multiple Comparison Test); # # or ** p<0.001, # or * p<0.05.

The effect of hypoxia on cell division was then examined using CellTrace staining. As expected with hypoxia alone, we see significant reductions in the estimated number of divisions in MOLM-13 and MV-4-11s, and a trending decrease in HL-60s (***Fig. 1C***). A decrease in cell divisions was also observed in normoxia upon DAC treatment in MOLM-13 and MV-4-11 cells, and AZA treatment in HL-60. Importantly, the introduction of HMAs in hypoxia showed no significant changes compared to the respective untreated (UT) cells. These results also support the overall growth trends seen in ***Fig. 1A***, and strong positive correlations were observed between the total viable cells and the estimated number of cell divisions (***Fig. 1C, Supplementary figure 2***). Here, we demonstrate that hypoxia significantly reduces growth in AML cell lines, and that the reductions associated with cell division may have implications on HMA incorporation.

### Hypomethylation induced by AZA is impaired in hypoxia

Given that HMAs are incorporated into DNA during replication [17-19], the reduced growth of AML cells observed in hypoxia was expected to impact treatment-induced hypomethylation. To investigate this, global DNA methylation levels were analysed using a post-bisulphite adaptor tagging (PBAT) and sequencing approach. In both normoxic and hypoxic conditions, HMA treatment significantly decreased DNA methylation levels in all cell lines relative to their respective UT controls (***Fig. 2***). However, when comparing the methylation levels of respective treatment groups between normoxic and hypoxic conditions, we see a significant difference in HMA efficacy. Particularly, AZA treatment was significantly less effective in hypoxia, with methylation levels being higher in all cell lines compared to treatment in normoxia. There was also a modest retention of DNA methylation observed when HL-60 cells were treated with DAC in hypoxia. In contrast, hypoxia showed no significant implications on DAC treatment in MOLM-13 and MV-4-11 cell lines. While surprising, these results highlight important differences between DAC and AZA, and suggest that DAC may maintain efficacy in hypoxia despite the suppressed growth of AML cells (***Fig. 1***).

**Figure 2:**
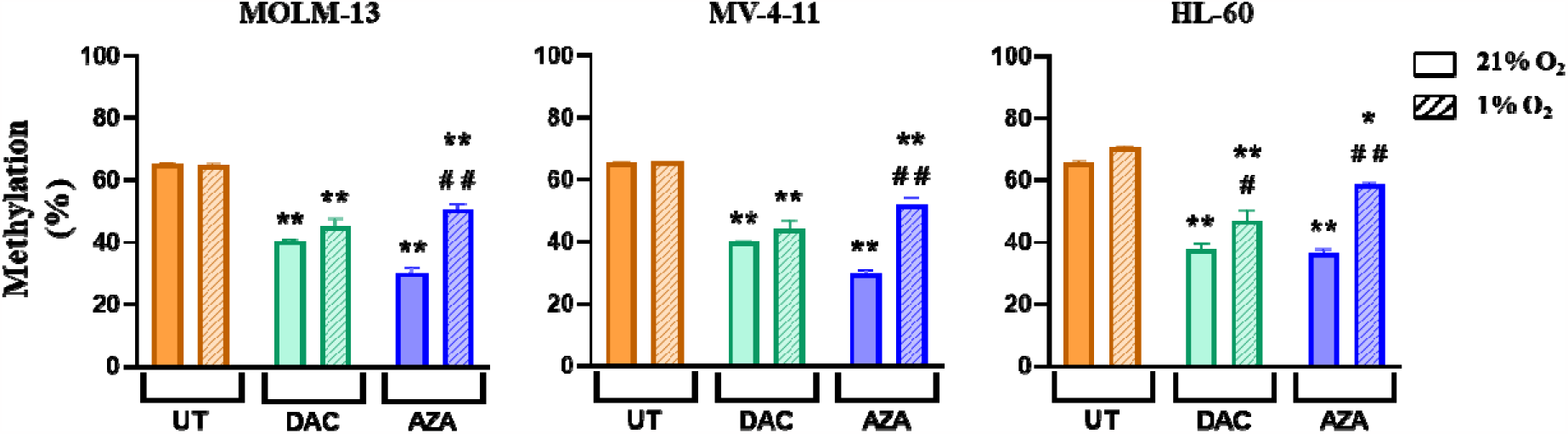
*The effect of hypoxia on HMA-induced DNA hypomethylation in AML cell lines*. DNA methylation from HMA-treated AML cell lines (MOLM-13, MV-4-11 and HL60) in normoxic (21% O_2_; solid) and hypoxic (1% O_2_; striped) conditions were examined using post-bisulphite adaptor tagging (PBAT) analysis. Data is presented as mean ± standard error of the mean for replicate experiments (MOLM-13 n=10; MV-4-11 n= 6; HL-60 n=6). # indicates significant difference between oxygen tensions within each treatment group. * indicates significant difference of treatment from the UT control within each oxygen tension. Two-way ANOVA (Tukey’s Multiple Comparison Test); # # or ** p<0.001, # or * p<0.05.

### Transcriptional responses typically induced by HMA treatment are altered in hypoxia

To explore the transcriptional implications of altered hypomethylation in hypoxia (***Fig. 2***), RNA-sequencing was performed in MOLM-13 cells. We identified a total of 4302 differentially expressed genes (DEGs) with DAC treatment (***Supplementary Table 2***) and 3849 DEGs with AZA (***Supplementary Table 3***). Hierarchical clustering was then performed to visualize gene expression changes within each treatment group. Here we see that treatment with AZA (***Fig. 3A***) and DAC (***Fig. 3B***) cause gene expression changes that are altered in hypoxia.

**Figure 3:**
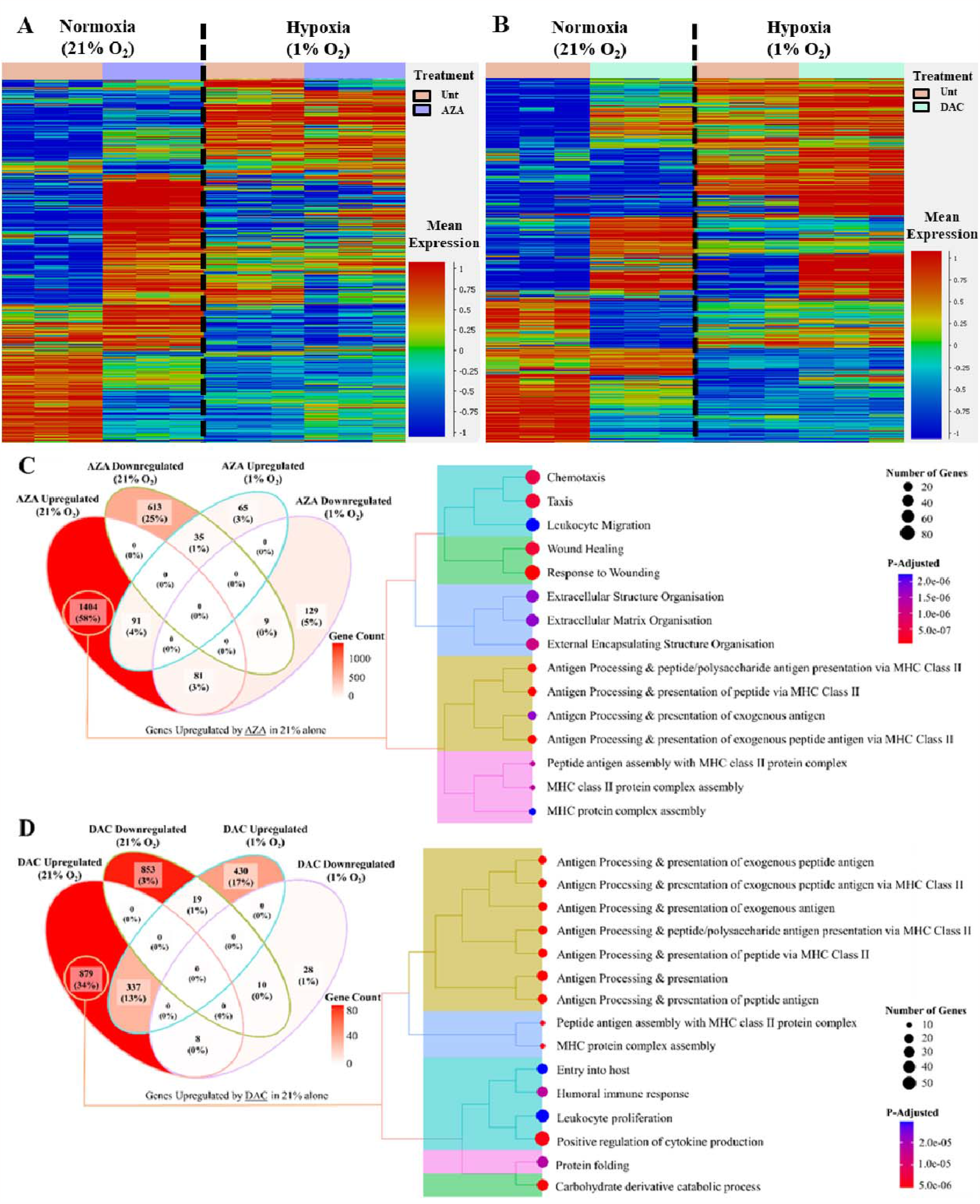
*The effect of hypoxia on HMA-induced transcriptional changes in MOLM-13*. RNA sequencing was performed on MOLM-13 cells to generate a heatmap of differentially expressed genes (DEGs) comparing the effect of (**A**) AZA, or (**B**) DAC within each oxygen tension (DESeq2; p≤0.01 and fold change >1, or <-1). (**C**) depicts a venn diagram summarizing genes up- or down-regulated by AZA in normoxic (21%) or hypoxic (1%) conditions, while **(D)** summarizes genes up- or down-regulated by DAC. Venn diagrams are accompanied by treeplots that present the top 15 gene pathways upregulated by AZA or DAC in 21% O_2_ alone. Pathways were selected using the clusterProfiler program in RStudio comparing against a human reference genome (org.Hs.eg.db). Pathways were determined using an FDR p-adjustment method, and a p-value cutoff of 0.05, and a q-value cutoff 0.2. of

We next wished to identify genes that were uniquely upregulated following HMA treatment in normoxia to highlight the inhibitory effect that may be imposed by hypoxia. As such, DEG lists were compared (***Fig. 3C, D***). Of the 2017 genes (83%) that were uniquely altered in normoxia following AZA treatment, 1404 genes (58%) were up-regulated. Gene ontology analysis revealed that these genes showed an over-representation of processes related to chemotaxis, wound healing, extracellular matrix organisation and antigen presentation (***Fig. 3C, Supplementary Table 4***). With DAC treatment, 1732 genes (67%) were uniquely altered in normoxia, with 879 (34%) of these being up-regulated and enriched for antigen presentation, cytokine production, protein folding and carbohydrate metabolism (***Fig. 3D, Supplementary Table 5***). These results highlight further differences between DAC and AZA as individual treatments, but also indicate that some transcriptional responses induced by HMAs in normoxia (e.g., antigen presentation), are not observed in hypoxia. This could have important implications for downstream response to HMAs in hypoxic environments, like the bone marrow.

### Hypoxia impairs HMA-induced HLA-DR expression

Antigen presentation is a critical pathway required for the detection and elimination of pathogenic or abnormal cells by activating the innate or adaptive immune system [47]. Within the antigen presentation pathways up-regulated by HMAs in normoxia alone (***Fig. 3***.), a striking number of the DEGs were related to human leukocyte antigen (HLA) genes (***Supplementary Table 4, 5***) – which encode proteins that help the immune system identify cells as “self” or “non-self” [48]. The numerous members of this family [48] are divided into class I and class II gene isotypes, known for activating CD8+ T cells/natural killer cells or CD4+ T cells, respectively [48]. Importantly, however, the most notable upregulation was observed with class II isotypes following HMA treatment in normoxia (***Fig. 4A, Supplementary Table 6, 7***), suggesting that HMAs may be unable to promote a class II immune response in hypoxia.

**Figure 4.**
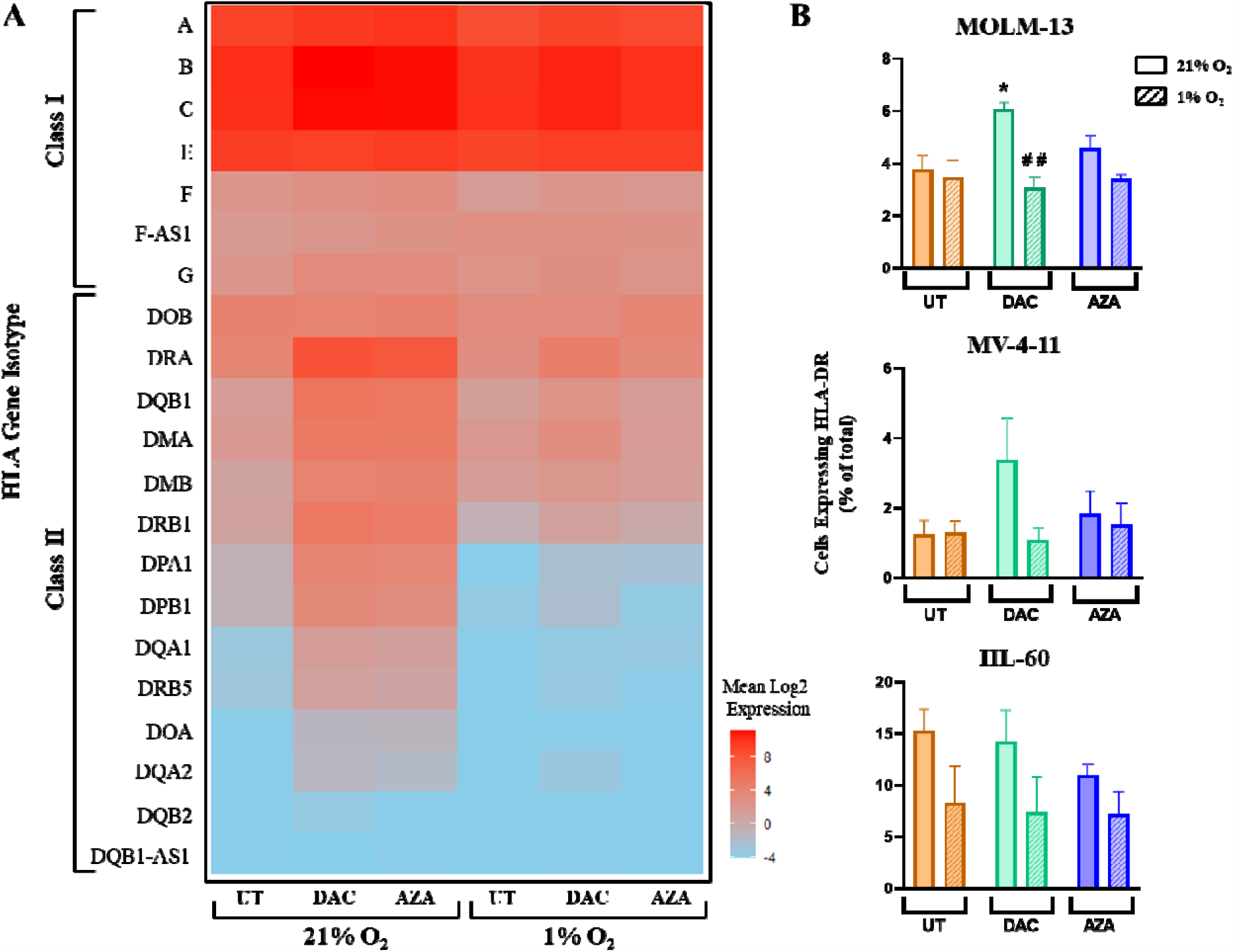
*The effect of hypoxia on HMA-induced HLA expression*. (**A**) MOLM-13 RNA-seq data was used to analyse the log2 normalized read counts for HLA transcripts following treatment (Untreated – UT, Decitabine – DAC, Azacitidine – AZA) in either normoxic (21% O_2_) or hypoxic (1% O_2_) conditions (n=3). HLA isotypes were grouped by class and manually ordered by expression level. (**B**) HLA-DR cell surface protein expression in MOLM-13, MV-4-11, and HL-60s following HMA treatment (UT, DAC, AZA) in normoxic (21% O_2_; unshaded) or hypoxic (1% O_2_; shaded) conditions. Data is presented as mean ± standard error of the mean for replicate experiments (MOLM-13 n=3; MV-4-11 n= 3; HL-60 n=5). # indicates significant difference between oxygen tensions within each treatment group. * indicates significant difference of treatment from UT control within each oxygen tension. One-way ANOVA (Tukey’s multiple comparisons); # # or ** p<0.001, # or * p<0.05.

To validate these observations, flow cytometry was used to assess HLA class I (HLA-G) and HLA class II (HLA-DR, -DP, and -DQ) cell-surface protein expression levels in all cell lines (***Fig 4B***., ***Supplementary Figure 3A, B***). In normoxic conditions, DAC treatment either significantly increased (e.g. MOLM-13), or trended towards increasing (e.g. MV-4-11), the proportion of cells with membrane expression of HLA-DR a key cell surface molecule responsible for initiating the inflammatory response via antigen presentation [49]. Importantly, when cells were treated with HMAs in hypoxia, HLA-DR expression was unchanged in all cell lines respective to their untreated controls. Similar results were detected using the polyclonal HLA-DR, -DP and -DQ antibody, and monoclonal HLA-G antibody, where minimal changes were observed in hypoxia (***Supplementary Figure 3A, B***). These results support the idea that hypoxia may impair HMA-induced antigen presentation.

## 4. Discussion

AML cells reside in the bone marrow microenvironment where low oxygen tensions may offer protection from anti-cancer agents. In this study we investigated the direct impact of hypoxia on HMA treatment response in AML cell lines. Consistent with previous reports [28, 50, 51], our results reflected significant reductions in cell growth during hypoxia (***Fig. 1A***). When HMAs were added in these hypoxic conditions, no additional effects on cell growth were observed. Since HMAs are primarily replication-dependent therapies [19, 52], the growth reductions seen in hypoxia suggests implications for their overall efficacy.

Direct examination of global HMA-induced hypomethylation in hypoxia revealed surprising differences between DAC and AZA. While DAC was capable of reducing DNA methylation to a similar extent in both oxygen tensions, the efficacy of AZA was significantly hindered in the context of hypoxia (***Fig. 2***). These observations are consistent with a previous study of MDS patients, where AZA non-responders had a higher proportion of quiescent LSCs in the hypoxic bone marrow [35]. AZA may be less effective than DAC in hypoxia because, unlike DAC, AZA primarily incorporates into RNA with only 10-35% of AZA incorporated into DNA [19, 53]. We speculate that the reduced rate of DNA replication in hypoxia could favour RNA incorporation of AZA. This would reduce the amount of AZA available for DNA incorporation and could explain the impaired hypomethylation observed following AZA treatment in hypoxia.

Since DNA methylation plays direct roles in gene expression, HMA-induced changes to the epigenome, particularly in cancers, can reactivate tumour suppressor genes [54-57]. As DAC and AZA present differences in their ability to hypomethylate the genome in hypoxia, RNA sequencing was used to explore common and unique downstream pathways that may be impacted (***Fig. 3A, 3B***). Interestingly, the majority of AZA-induced gene expression changes were observed in normoxia only (83% of AZA DEGs; ***Fig. 3C***), consistent with the impaired hypomethylation observed for AZA treatment in hypoxia (***Fig. 2***). In contrast, relatively few DAC-induced transcriptional changes were unique to normoxia (37% of DAC DEGs), demonstrating that some DAC-induced transcriptional changes are observed in both oxygen tensions. Despite these differences in transcriptional response following AZA and DAC treatment in hypoxia, we found important commonalities between treatments. Specifically, genes involved in antigen presentation were uniquely up-regulated following both HMA treatments in normoxia, but not hypoxia (**Fig. 3C, 3D**). This suggests that the HMA-induced immune response may be suppressed in hypoxic conditions.

Importantly, genes affecting human leukocyte antigens (HLA), which are critical components for antigen presentation and initiation of the immune response [58, 59], were among the genes up-regulated by HMAs in normoxia only. HLAs are immune glycoproteins divided into two main classes. HLA class I proteins activate our innate and adaptive immunity through CD8+ T-cells and natural killer cells, respectively. In contrast, class II proteins only contribute to our adaptive immunity by activating regulatory CD4+ T-cells [58, 59]. Our results demonstrate that, while HLA class I gene expression appears unaffected by HMA treatment in both oxygen conditions, class II genes are uniquely up-regulated in normoxia following HMA treament, with no notable changes observed in hypoxia (***Fig. 4A***). This observation was confirmed at the protein level for the class II antigen, HLA-DR (***Fig. 4B***). Previous studies have shown that expression of class II HLAs [60] can be increased by HMA treatment in AML [61-63], but our study is the first to show that this response may be impaired by hypoxic conditions. Consistent with reports that LSCs residing in the hypoxic bone marrow can present increased immune evasion [64-66] and chemoresistance [67-70], our study suggests that quiescent cells in the hypoxic BM, may be able to avoid HMA-induced expression of antigen presentation-related genes. This could allow them to escape immune surveillance and retain the long-term repopulation capacity required for relapse.

## 5. Conclusions

Collectively, this study provides insight into the importance of BM microenvironmental factors for AML treatment efficacy. Not only does hypoxia allow AML cells to enter a quiescent-like state, but it also decreases the efficacy of HMAs, altering downstream transcriptional responses. As a result, alternative treatment strategies that can re-initiate the cell cycle in LSCs or target their immune pathways may be required to enhance HMA treatment response in hypoxic conditions. Future studies should examine the effect of HMAs and combinatorial treatments on cancer cells in various tissues (e.g., the peripheral blood and bone marrow) to ensure consistent therapeutic efficacy throughout the body.

## 6. Declarations

### Ethics Approval and Consent to Participate

Not applicable.

### Consent for Publication

Not applicable.

### Availability of Data and Materials

Sequencing data will be made available to reviewers upon request and published upon manuscript acceptance.

### Author’s Contributions

S.H. performed experiments and data analysis, prepared figures, and drafted the manuscript. S.M.B. processed sequencing data and supervised data analysis. S.K. and D.R.B. supervised experiments.

H.J.L. conceived and oversaw the project and acquired funding. S.M.B., D.R.B. and H.J.L edited the manuscript. All authors reviewed the manuscript and approved the final version. Correspondence should be addressed to H.J.L.

## Acknowledgements

The authors thank Dr. Heather Murray from the University of Newcastle for providing a critical review of the manuscript.

