## Supplementary Figure for "Hypoxia alters the effects of hypomethylating agents in acute myeloid leukaemia cells"

### MOLM-13

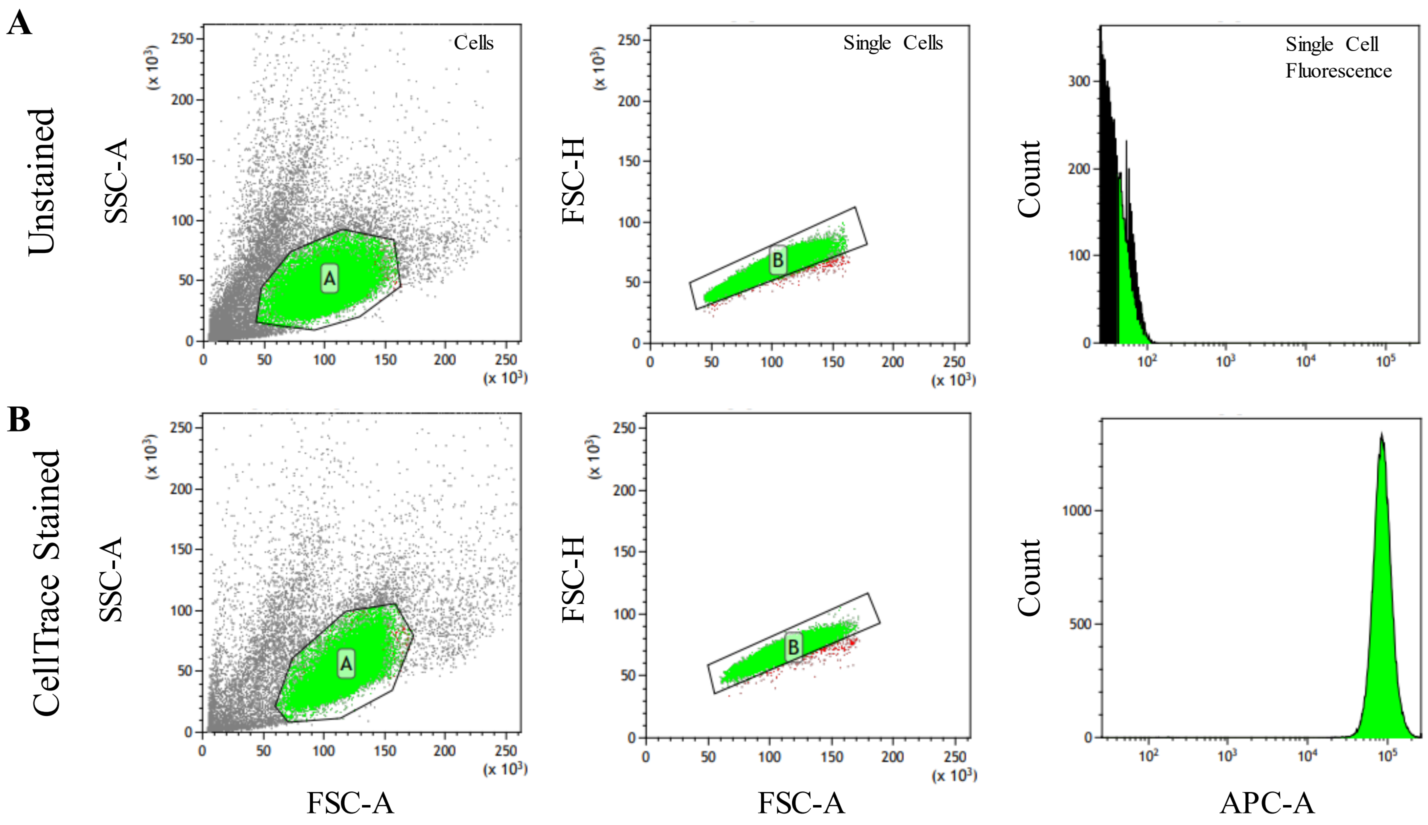


**Supplementary Figure 1:** *Gating strategies for detecting CellTrace fluorescence in singlet cells.* MOLM-13 cells were pre-conditioned to normoxic conditions for 72 hours (Day 0) before staining with CellTrace Far Red Cell Proliferation Stain (APC) and subsequent treatment with HMAs in normoxic and hypoxic conditions. After pre-conditioning, a 'Day 0' negative (unstained; ***A***) and positive (stained; ***B***) control were examined using the FACSCanto II flow cytometer. Data was imported into Kaluza software and singlet cells were analysed using the gating strategies depicted in the representative images shown. The single cell fluorescence illustrates the number (count) of gated cells with the bound fluorescent antibody. Side Scatter-Area (SSC-A); Forward Scatter-Area (FSC-A); Forward Scatter-Height (FSC-H); Allophycocyanin-Area (APC-A).

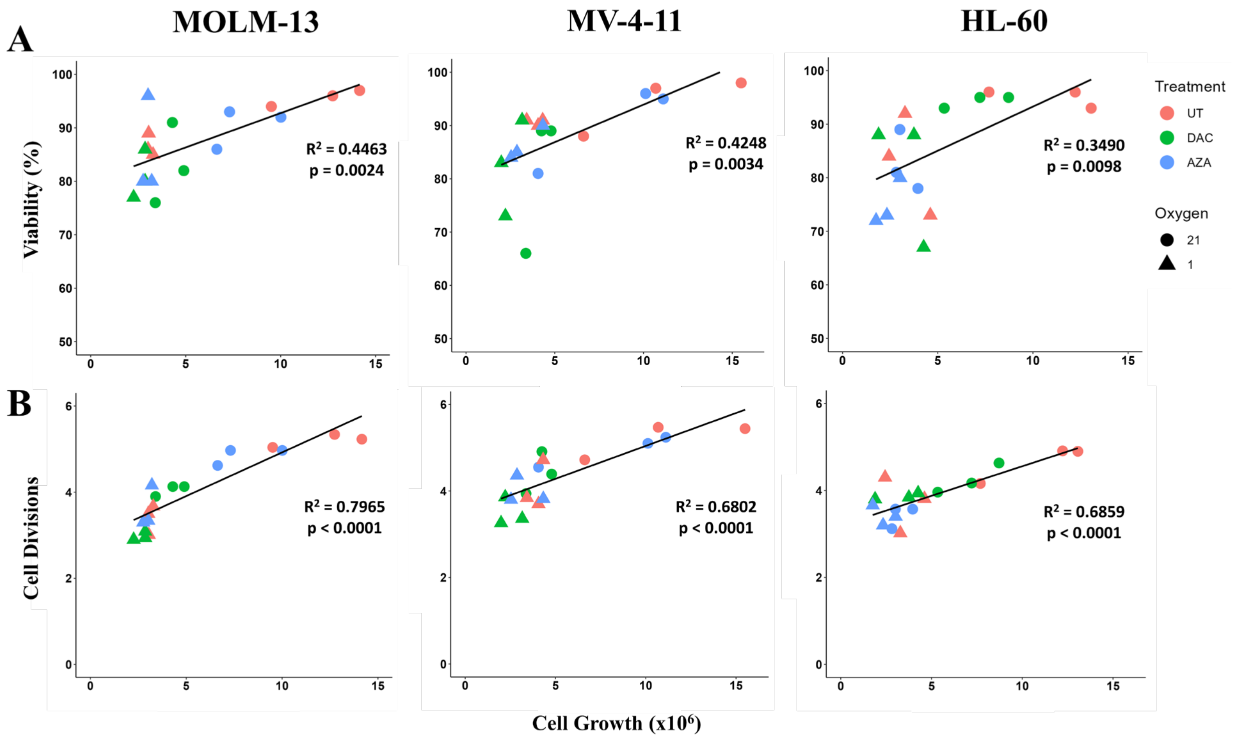


**Supplementary Figure 2**. *Correlations between cell growth and viability, or estimated cell divisions.* Cell growth values for each AML cell line were correlated to either the viability **(A)** or estimated cell divisions **(B)** of the same sample. Untreated (UT, Pink), Decitabine (DAC; Green), and Azacitidine (AZA; Blue) treated cells were allocated based on treatment in 21% (circle) or 1% (triangle) $O_2$. Pearson's correlation coefficient ($R^2$) and significance were generated using GraphPad software. Significance was calculated using a two-tailed paired t-test (95% CI).

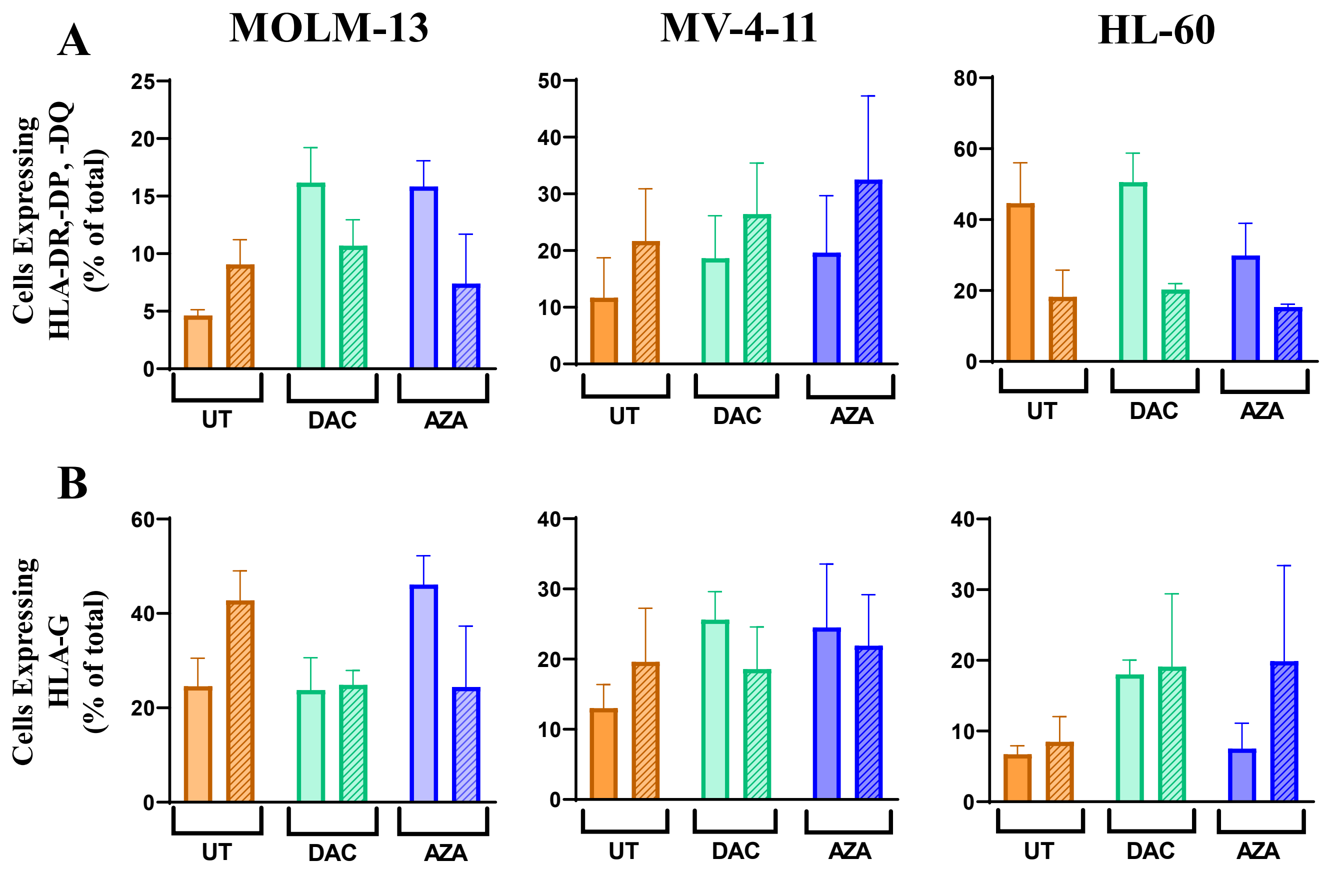


**Supplementary Figure 3**. *The effect of hypoxia on HMA-induced protein expression of HLA-DR, -DP, -DQ, and -G* . ***(A)*** Polyclonal surface protein expression of HLA-DR, -DP and –DQ, or **(B)** monoclonal expression of HLA-G, in MOLM-13, MV-4-11, and HL-60s following HMA treatment (UT, DAC, AZA) in normoxic (21% $O_2$; unshaded) or hypoxic (1% $O_2$; shaded) conditions. One-way ANOVA (Tukey's multiple comparisons) was used to compare effects between 21% $O_2$ vs. 1% $O_2$ of each HMA treatment group (#) or HMA vs UT within each oxygen tension (*); # # or ** $p<0.001$, # or * $p<0.05$. Data was  presented as a mean ± standard error of the mean.
